# Identifying deleterious noncoding variation through gain and loss of CTCF binding activity

**DOI:** 10.1101/2024.09.04.609712

**Authors:** Colby Tubbs, Mary Lauren Benton, Evonne McArthur, John A. Capra, Douglas M. Ruderfer

**Affiliations:** Division of Genetic Medicine, Department of Medicine, Vanderbilt Genetics Institute, Vanderbilt University Medical Center, Nashville, TN; Department of Computer Science, Baylor University, TX; Department of Medicine, University of Washington, Seattle, WA; Department of Epidemiology and Biostatistics, Bakar Computational Health Sciences Institute, University of California at San Francisco, CA; Center for Digital Genomic Medicine, Vanderbilt University Medical Center, Nashville, TN

## Abstract

Noncoding single nucleotide variants are the predominant class of genetic variation in whole genome sequencing and are key drivers of phenotypic variation. However, their functional annotation remains challenging. To address this, we develop a hypothesis-driven functional annotation scheme for CTCF binding sites given CTCF’s critical roles in gene regulation and extensive profiling in regulatory datasets. We synthesize CTCF’s binding patterns at 1,063,879 genomic loci across 214 biological contexts into a summary metric, which we refer to as binding activity. We find that binding activity is significantly enriched for both conserved nucleotides (Pearson R = 0.31, p < 2.2 x 10^-16^) and sequences that contain high-quality CTCF binding motifs (Pearson R = 0.63, p = 2.9 x 10^-12^). We then integrate binding activity with high confidence change in precision weight matrix scores. By applying this framework to 1,253,330 SNVs in gnomAD, we explore signatures of selection acting against the disruption of CTCF binding. We find a strong, positive relationship between the mutability adjusted proportion of singletons (MAPS) metric and the loss of CTCF binding at loci with high *in vitro* activity (Pearson R = 0.67, p = 1.5 x 10^-14^). To contextualize these findings, we apply MAPS to other functional classes of variation and find that a subset of 198,149 loss of CTCF binding variants are observed as infrequently as missense variants. This work implicates these thousands of rare, noncoding variants that disrupt CTCF binding for further functional studies while providing a blueprint for the interpretable annotation of noncoding variants.

## Introduction

Noncoding genetic variation functions as a major driver of phenotypic variation^1,2^; accounting for over 90% of GWAS loci^3,4^, explaining a substantial proportion of tissue-specific eQTLs^5,6^ and driving pathogenic gene expression in a growing number of rare disorders^7–10^. Despite their clear importance, extracting functional regulatory variants from the practically limitless nonfunctional mutations possible in noncoding DNA remains a major challenge. Whole-genome sequencing (WGS) captures almost entirely noncoding variation given that 1% of the genome is protein-coding^11^. As the field moves to these data ^12–14^ to expand genomic understanding beyond proteins, there is a critical need to understand the effects of noncoding variants and their contribution to traits and diseases. Developing approaches to functionally annotate noncoding variants is crucial to advancing our ability to prioritize among the expanding catalog of genetic variation implicated in human disease studies for further analysis.

The functional effects of a variant in coding sequence can be assessed as a product of the gene characteristics (e.g. constraint, haploinsufficient, etc.)^15,16^ and the impact the variant has on the protein sequence^17^ (e.g. nonsense, missense, etc.). Efforts to apply a similar approach to noncoding DNA is difficult for several reasons. First, there is no noncoding equivalent to the triplet amino acid code that governs coding sequence^18^. Second, the regulation of gene expression by *cis*-regulatory elements (CREs) is cell-type specific ^19–21^, which would require extensive high-throughput genomic profiling across hundreds or even thousands of biological contexts to comprehensively identify. Third, CRE annotations typically span hundreds of base pairs and the mapping between sequence and function is incompletely understood, making the identification of functional nucleotides difficult ^22,23^. Finally, the relationship between regulatory dysfunction and molecular phenotypes (i.e. gene expression) can be complex and indirect^24^.

Given the many challenges of functionally annotating noncoding variants, existing methods often rely on hypothesis-free approaches by leveraging evolutionary sequence conservation or machine learning algorithms that lack specific insights into the molecular functions of the variants prioritized. In these paradigms, functional constraint on individual nucleotides is quantified by aligning reference sequences from closely related species^25–28^. Single-nucleotide variants (SNVs) that occur at positions that are depleted of interspecies substitutions are considered constrained and are interpreted as more likely to contribute to phenotypes. Alternatively, machine learning algorithms are used to summarize a variant’s relationship with numerous and diverse functional genomic annotations into a single pathogenicity score^29–33^. While powerful, conservation metrics and machine learning algorithms can mask intuitive insight into their output scores and often require extensive follow-up to understand a variant’s biological function.

Several concurrent developments provide opportunities to design methods for the interpretation of noncoding variants that provide mechanistic insights into function akin to approaches for coding sequence. First, there has been extensive development in the systematic identification and characterization of regulatory DNA sequences. The ENCODE consortium has now released detailed, integrated maps of candidate cis-regulatory element (cCRE) activity across thousands of biological contexts^23,34^. These annotations measure DNA accessibility and transcription factor (TF) binding at millions of loci through DNase-Seq and ChIP-Seq. Second, new experimental and computational methods have enabled the quantification of TF-DNA binding preferences, which can be represented in precision-weight matrices (PWMs). PWMs quantify the contribution of DNA sequence to binding energy at base pair resolution and are now widely available for hundreds of TFs^35,36^. The combination of regulatory sequences and variant level effects on those sequences provides an opportunity to improve interpretation of noncoding variants in a similar vein as interpretation of coding variants.

Here, we capitalize on this progress by developing a hypothesis-driven annotation scheme for SNVs that disrupt transcription factor binding sites (TFBSs). We focus our efforts on binding sites for CCCTC binding factor (CTCF) as an illustrative case study for two reasons: 1) CTCF’s crucial roles in regulating gene expression through 3D genome organization make its binding sites likely to harbor functional variation^37–41^ and 2) its extensive profiling in different cellular contexts allows for the effective annotation of its binding patterns across hundreds of cell and tissue contexts^22,42,43^. To annotate SNVs that disrupt CTCF binding sites (CBS), we integrated CTCF’s canonical binding motif, ENCODE’s cCREs and large-scale genetic variation data. We evaluated our approach by comparing our annotations to metrics of conservation, deleteriousness and evidence for negative selection in gnomAD, the largest publicly available WGS database.

We demonstrate a scalable blueprint for the functional annotation of noncoding SNVs and provide a comprehensive evaluation of constraint on variants that are predicted to alter CBS activity. We detect strong signatures of negative selection acting on thousands of rare variants predicted to induce the loss of CTCF binding at loci with high *in vitro* binding activity. This work explores a critical step in interpreting noncoding variation and will enable improved prioritization of noncoding variants for future disease studies.

## Results

### 2.1. Framework for the functional annotation of CTCF binding site variants

ChIP-Seq captures TF binding but reports regions that span hundreds of basepairs and provides little means to interpret SNVs^44^. Conversely, PWMs can capture variant-level contributions to binding energy but have enormous false-positive rates^45^. Here, we build an approach that combines ENCODE’s cCRE framework with binding information represented in CTCF’s PWM **(Figure 1)**. The cCREs provide estimates of the strength of CTCF binding at millions of representative DNase hypersensitivity regions (rDHSs) in the genome and reports this signal as a Z-score relative to all loci tested in each cell line or tissue sample (henceforth referred to as biosamples, **Figure 1A**). We extracted CTCF’s ChIP-Seq Z-score at 1,063,879 rDHSs across 214 biosamples. These rDHSs were typically several hundred basepairs in length (mean = 273bp, SD = 67bp). To interpret SNVs within these regions, we reduced each rDHS to any and all subsequences matching CTCF’s binding motif **(Methods)**. Of all CTCF motif matches in the genome (**Supplementary Figure 1A**), we identified 355,418 that are overlapped by a rDHS, which we refer to simply as CBSs **(Supplementary Table 1).**

**Figure 1.**
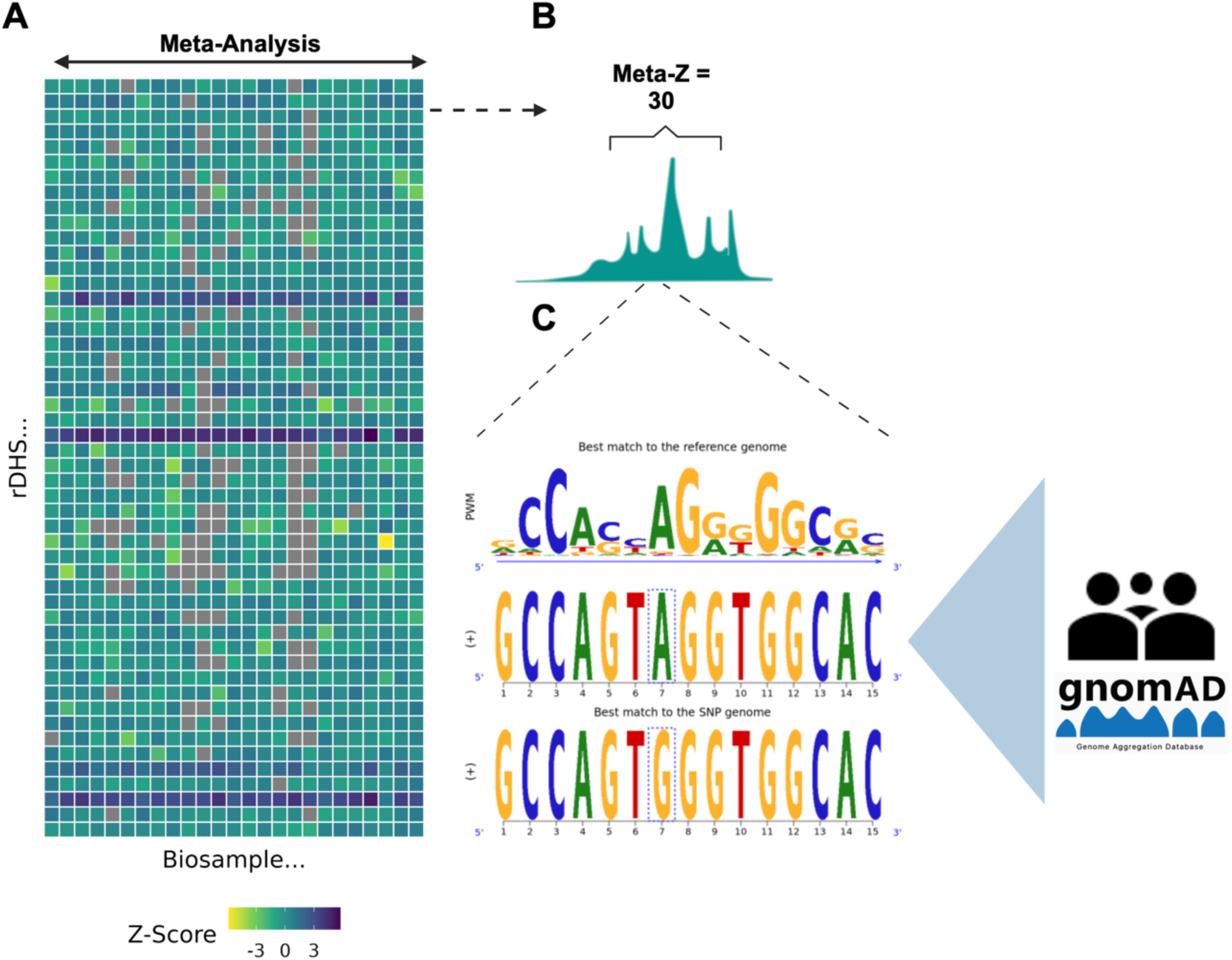
Schematic overview of the study approach. **A)** Heatmap of CTCF’s ChIP-Seq binding activity Z-score at selected rDHS and biosamples. The X-axis represents 25 of 214 randomly sampled biosamples and the Y-axis represents 100 of 1,063,879 randomly sampled rDHS. Coloring corresponds to the strength of binding (Z-score) for a given biosample and rDHS. **B)** Schematic illustration of a theoretical distribution of binding activity Z-scores for a single rDHS across 25 biosamples. We utilize these data to develop a novel annotation for CTCF’s observed binding patterns across these biosamples, which we refer to as binding activity. Here, binding activity would be quantified as a single summary score by meta-analyzing the depicted distribution of Z-scores using Stouffer’s Method. **C)** Schematic illustrating the integration of binding activity with the functional annotation of SNVs through ΔPWM scores. Pictured as an example is a putative loss of binding SNV that has reduced binding energy in the alternate allele. We calculate high confidence ΔPWM scores for all candidate SNVs in 76k WGS samples from gnomAD.

### 2.2. Developing a summary annotation of CTCF’s binding activity *in vitro*

PWMs measure binding using sequence content only and do not capture the multitude of extrinsic factors that govern when TFs bind DNA. This property of PWMs imparts a high false positive rate in that most motifs they detect will never function in binding TFs *in vitro* (i.e. the Motif Futility Theorem^45^). Here, we provide direct support for this concept; most rDHS (73%) did not contain an underlying CTCF PWM match, regardless of their cCRE classification (**Supplementary Figure 1B**). Of the rDHSs that did contain one or more motifs (n= 290,793), most contained fewer than 4 (**Supplementary Figure 1C**). We next evaluated how frequently a CTCF bound rDHS is replicated in at least more than one biosample. We found that at the minimal threshold used to define CTCF binding in the ENCODE framework (Z-score > 1.64), a substantial proportion of binding events were observed in only a single biosample (**Supplementary Figure 1D**). These data suggests that either modality (PWM or ChIP-Seq) alone will be insufficient for the robust detection of active CTCF binding sequence.

We thus sought to quantify the strength and replication of CTCF binding at each CBS into a single metric (i.e. an intolerance metric for a gene^15,16,46^). CTCF’s binding activity was summarized at each rDHS by meta-analyzing its distribution of binding activity Z-scores across all biosamples (mean = 2.09, SD= 11.09, **Figure 2A**, **Methods**). For simplicity, we refer to this annotation as binding activity.

**Figure 2.**
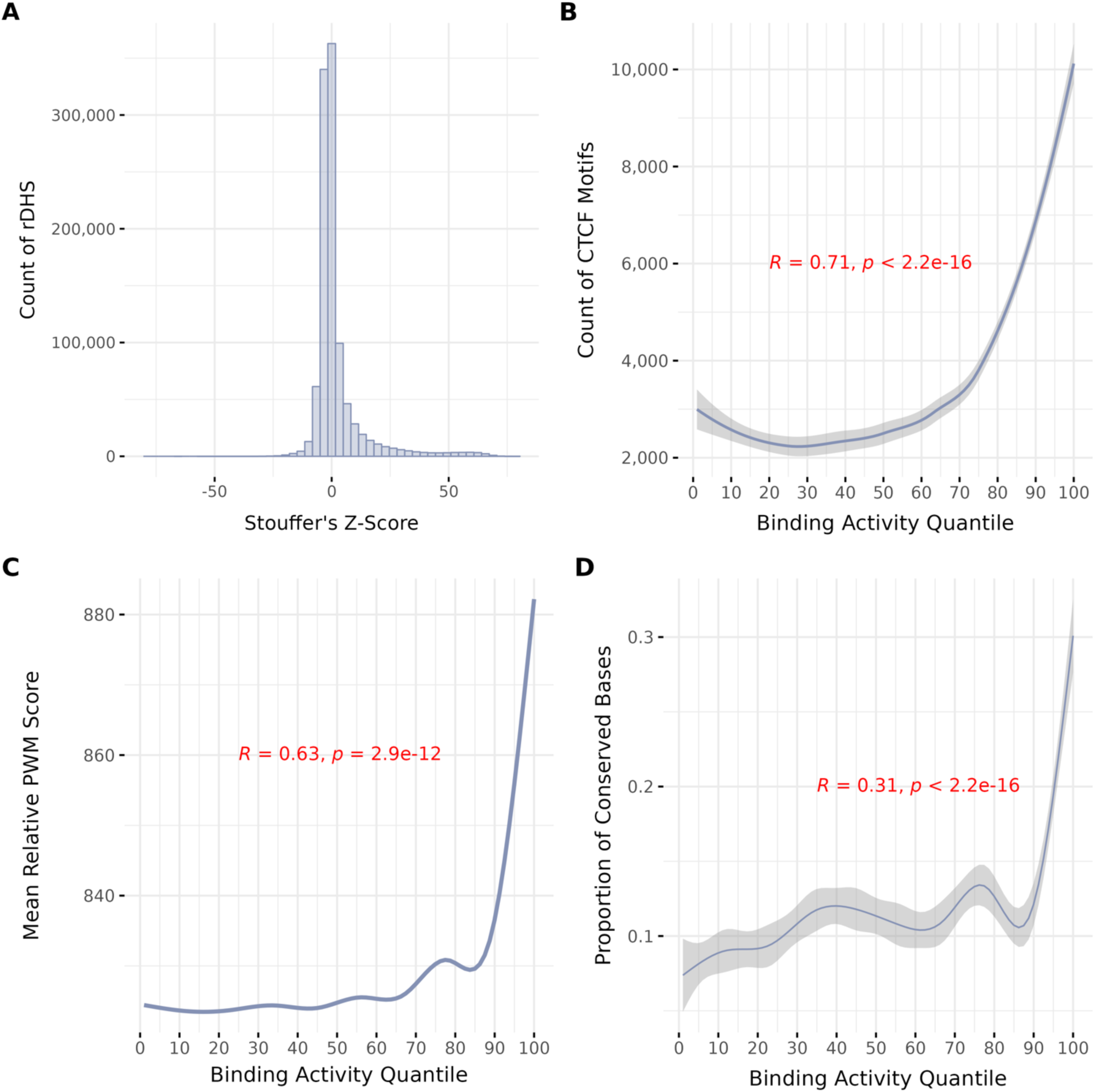
CTCF binding activity identifies sequence enriched for functional signatures. **A)** Distribution of meta-analyzed Z-scores for all 1,063,879 rDHS used in this study. Meta-analysis was conducted using Stouffer’s Method. **B)** The frequency of overlap between high-quality CTCF motifs and all rDHS belonging to a given quantile of binding activity. Quantiles were created by placing meta-analyzed Z-scores for all rDHS from A) into 100 equal sized bins. Pearson correlation was used for statistical evaluation. **C)** The mean PWM score for all CTCF motifs that overlap a rDHS in each quantile of binding activity. Pearson correlation was used for statistical evaluation. **D)** Relationship between evolutionary sequence conservation of CTCF motif sequence and binding activity quantiles. Conservation was measured as the proportion of conserved CTCF motif positions within each quantile using PhlyoP100 scores. A score threshold of 1 was used to define conserved positions. Pearson correlation was used for statistical evaluation.

To validate the use of binding activity in differentiating CBS with robust activity, we quantified its relationship with characteristics that are likely to signal authentic binding sequences. We note that we evaluate binding activity by summarizing characteristics across its distribution, in the form of quantiles **(Methods, Supplementary Figure 2)**. First, we observed an enrichment of CBSs (**Figure 2B**) occurring in high quantiles of binding activity (Pearson R = 0.71, p < 2.2 x 10^-16^). We then quantified each CBS’s predicted binding affinity for CTCF through its PWM **(Methods)** and observed a strong correlation between binding activity quantiles and predicted binding energy (Pearson R = 0.63, p = 2.9 x 10^-12^, **Figure 2C)**. Second, we evaluated the relationship between binding activity quantiles and functional constraint by annotating greater than 99% of the 5.3 Mb of CTCF binding sequence with evolutionary conservation metrics. PhlyoP scores quantify conservation at single nucleotides, making them well-suited for assessing constraint at high resolution. We calculated a proportion of conserved PhlyoP scores for each binding activity quantile and observed significantly higher proportions of conserved positions belonging to motifs in higher binding activity quantiles (Pearson R = 0.31, p < 2.2 x 10^-16^, **Figure 2D)**. Although conservation scores vary in their methodology, this correlation was consistent across multiple metrics (**Supplementary Figure 3**).

### 2.3. Integrating binding activity with variant-level functional annotation in gnomAD

Next, we sought to implement a functional annotation scheme for SNVs by integrating binding activity with the information encoded by PWMs. First, we identified all SNVs in gnomAD that disrupt a CBS (n=1,253,330, **Methods**). For each SNV in our dataset, we calculated its change from reference PWM (ΔPWM) and classified it as putative gain or loss of binding. We observed fewer gain of binding SNVs (n=228,470) compared to loss of binding SNVs (n=974,196) **(Figure 3A).** To assess confidence in each observed ΔPWM score, we calculated each score’s test statistic by implementing the importance sampling procedure developed by Zuo et al. 2015^47^ (**Methods**). We identified high confidence gain of binding (N=61,854) and loss of binding SNVs (N=504,507) using a significance threshold of p < 0.05 **(Figure 3B)**. We then integrated these predictions with binding activity by assigning each variant its motif’s activity score **(Figure 3C, Methods).** We observed significantly more high-confidence loss and gain of binding SNVs in high quantiles of binding activity (loss of binding Pearson R = 0.65, p = 2.1 x 10^-13^, gain of binding Pearson R = 0.67, p = 2.6 x 10^-14^).

**Figure 3.**
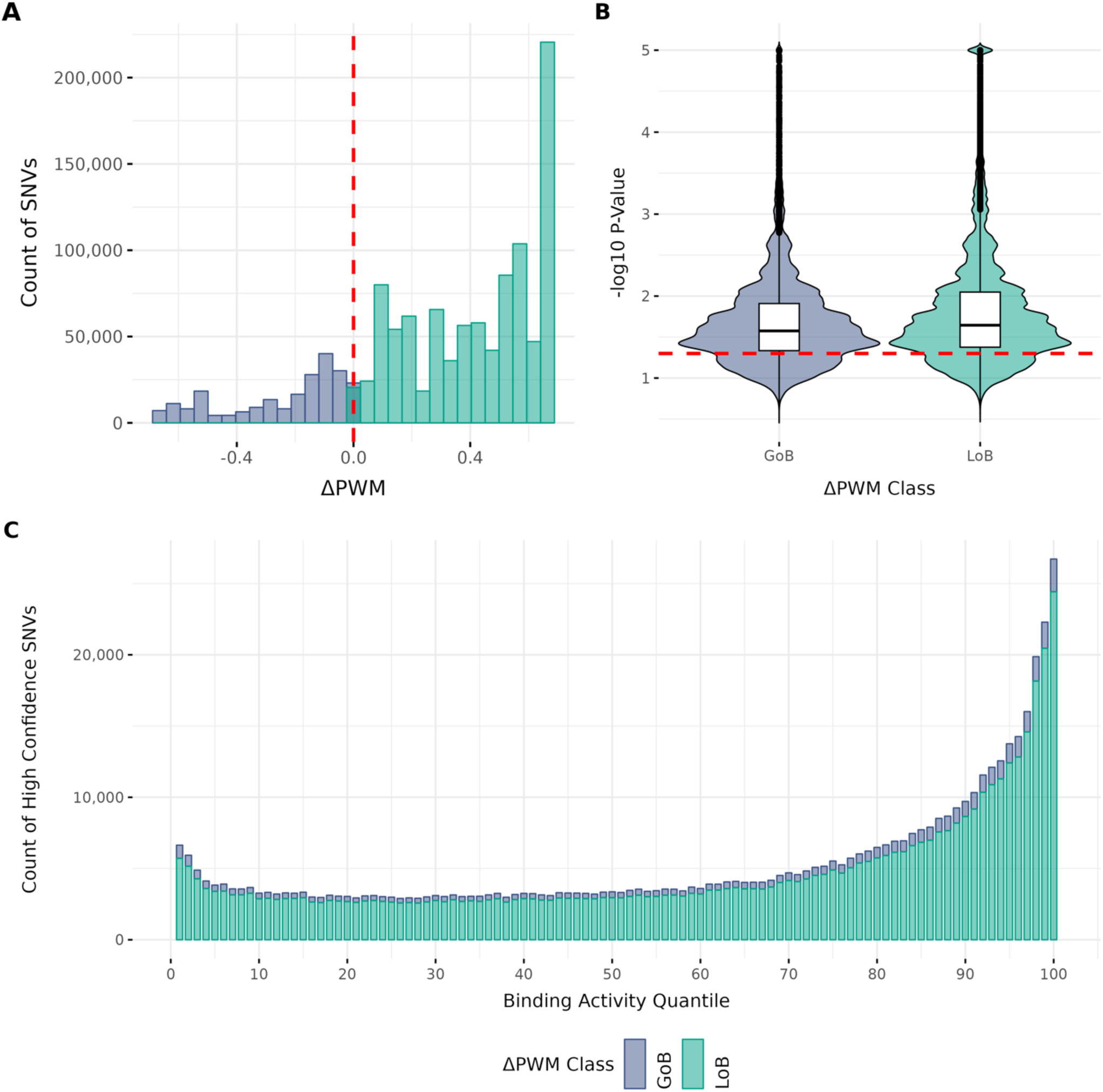
Functional annotation of CBS variants identifies primarily loss of binding in gnomAD. **A)** Distribution of ΔPWM scores for all SNVs in gnomAD that disrupt a CTCF motif with rDHS support. Gain of binding variants were considered as those that have a ΔPWM of <= 0. Loss of binding are defined as those having a ΔPWM of > 0. **B)** Distributions of ΔPWM test statistics for all SNVs in A) assessed through atSNP’s importance sampling procedure. Truncated p-values (approximately 0) were manually encoded as 10^-5^. Red line indicates the threshold used (p < 0.05) for distinguishing low and high confidence ΔPWM scores. **C)** Counts of high confidence ΔPWM scores in gnomAD for SNVs that disrupt a CTCF motif with rDHS support, binned by their binding activity quantile. Counts are stratified by their ΔPWM class (gain or loss of binding, GoB and LoB respectively).

### 2.4. Characterizing signatures of selection on the loss of CTCF binding activity

Deleterious genetic variants are purged from populations by natural selection, which enables the use of allele frequency as an informative metric for the utility of an annotation scheme in capturing functional variants^48,49^. For example, SNVs that disrupt genes are observed more rarely than those in intergenic space, while splicing SNVs are observed more rarely than those that fall in introns^11^. We evaluated our annotation approach in this framework by quantifying its relationship with different measures of selective constraint. To do this, we applied two measures: PHRED-scaled combined annotation dependent depletion (CADD)^50^ scores and the mutability adjusted proportion of singletons (MAPS) metric. First, we observed significantly higher proportions of predicted pathogenic variants (scaled CADD >= 10, Pearson R = 0.52, p < 2.2 x 10^-16^) for SNVs annotated as high-confidence loss of binding activity (**Figure 4A**). To assess the contribution of statistical testing on ΔPWM scores to this result, we calculated the proportion of predicted pathogenic variants for each binding activity quantile after stratifying variants by confidence in their ΔPWM score. At the 95^th^ quantile of activity or higher, we observed nominally higher frequencies of predicted pathogenic variants for high confidence ΔPWM scores compared to low confidence scores (high confidence ΔPWM mean proportion pathogenic=0.33, low confidence ΔPWM mean = 0.26, Student’s T-Test, p = 1.83 x 10^-2^, **Supplementary Figure 4A**). We observed no significant difference between low and high confidence ΔPWM scores for gain of binding SNVs in this analysis (**Supplementary Figure 4B**).

**Figure 4.**
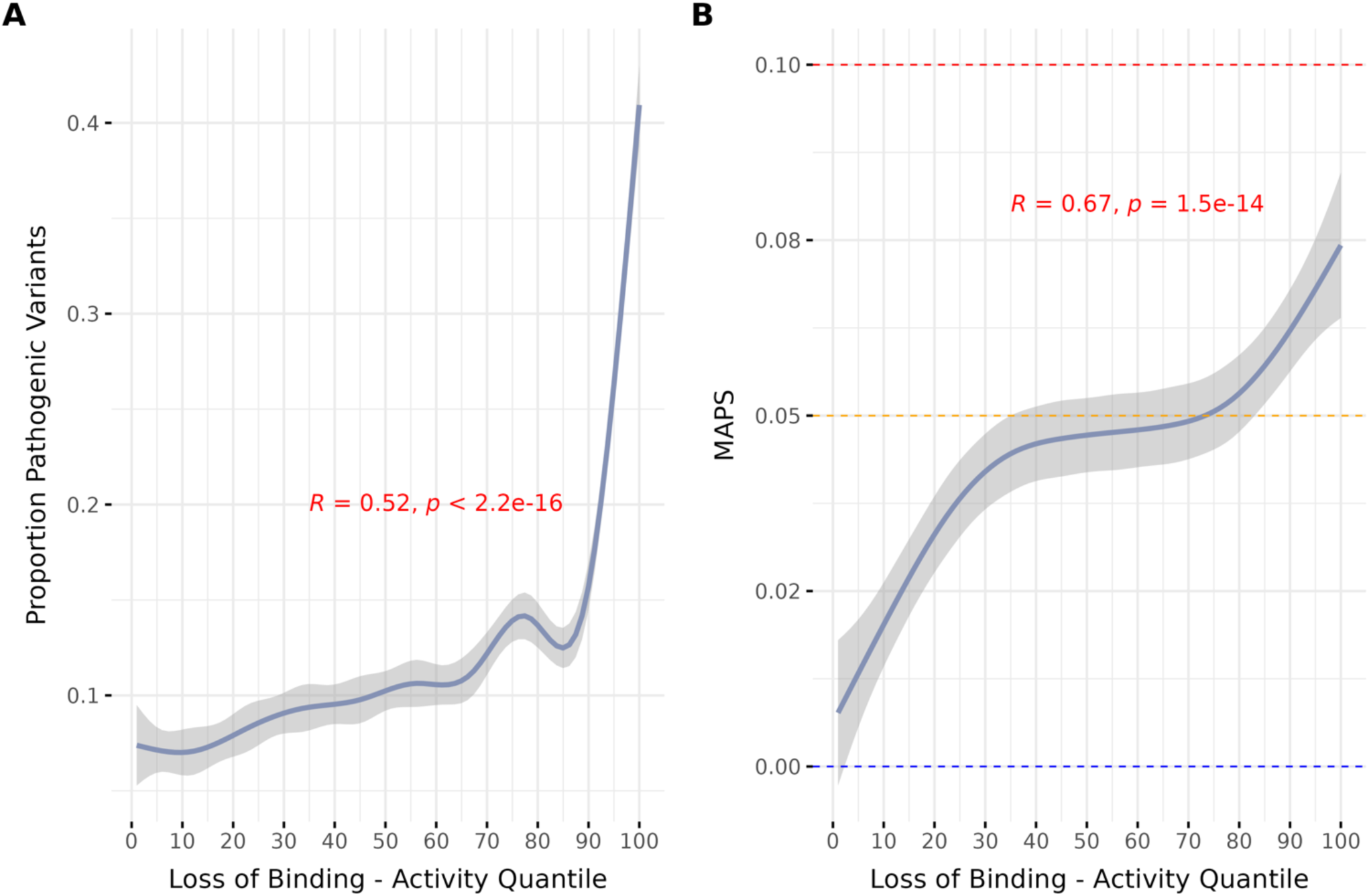
Loss of CTCF binding activity is under high levels of constraint in gnomAD. **A)** Relationship between proportion of putative pathogenic SNVs based on PHRED-scaled CADD scores and the loss of CTCF binding activity. A scaled CADD score of >=10 was used as a threshold of pathogenic or not. The Y-axis displays the proportion of pathogenic SNVs within each activity quantile. Error as a 95% bootstrapped confidence interval. **B)** Relationship between allele frequency as measured by MAPS and the loss of CTCF binding activity. For context, we display MAPS scores for synonymous (blue line), missense (orange) and splicing (red) variants in gnomAD.

Second, we calculated MAPS scores for all SNVs in each activity quantile, stratified by their predicted effect on binding affinity and the level of confidence in their ΔPWM scores. The loss of CTCF binding activity correlated significantly with MAPS, with a stronger overall effect for high-confidence SNVs (high-confidence Pearson R = 0.67, p = 1.5 x 10^-14^, **Figure 4B & Supplementary Figure 5A**). We note that while gain of binding at both high and low ΔPWM confidence displayed higher MAPS scores at higher levels of activity (**Supplementary Figure 5B**), there are substantially fewer SNVs in each bin. Consequently, we observe higher uncertainty on individual MAPS scores for both high and low confidence gain of binding SNVs (**Supplementary Figure 6**). To contextualize our findings, we calculated MAPS for different functional classes of intergenic and genic SNVs in gnomAD **(Supplementary Table 2)**. Loss of binding SNVs in binding motifs with low activity trended toward MAPS scores of likely neutral classes of variation (i.e., synonymous and intergenic), regardless of the confidence in the ΔPWM call. High-impact loss of binding variants (activity decile >= 90) had higher MAPS scores than missense variants (MAPS loss of binding = 0.07, MAPS missense = 0.04).

## Discussion

Expanded use of genome-sequencing combined with large-scale assays of regulatory function are providing novel opportunities for hypothesis driven functional annotation of noncoding SNVs. Here, we leveraged extensive regulatory data from ENCODE to develop a summary annotation for CTCF binding (binding activity) that effectively captures sequences enriched for functional constraint. We then integrated binding activity with high confidence ΔPWM scores to annotate over a million SNVs in gnomAD. We show that the loss of CTCF binding activity is associated with high levels of constraint and occurs infrequently in human populations. Together, these results provide evidence for an interpretable and novel annotation approach that prioritizes CTCF binding variants of likely functional consequence.

Understanding whether a CBS is functional is crucial for interpreting the consequence of genetic variants. We quantified the activity of a CBS by summarizing its strength and replication of binding across hundreds of biological contexts into a single statistic. This quantitative assessment of binding at a locus is conceptually appealing as it avoids reducing a complex interaction (TF-DNA binding) into a binary outcome. We further show its utility in several ways. First, we found that high binding activity correlates with functional signatures, including evolutionary conservation. This result was robust across multiple conservation metrics and suggests that binding activity effectively captures sequences that are likely to have functional consequences. Second, we observed that high confidence ΔPWM scores in motifs with low binding activity resemble likely neutral classes of variation (e.g. synonymous) in terms of their frequency in the population. Several databases have collectively assembled billions of ΔPWM scores by measuring SNVs against PWMs in an all-against-all framework^47,51^. Our results here suggest that a substantial proportion of these are likely false positives and that further annotation efforts should utilize a more targeted approach.

We observed strong signals of negative selection on SNVs that induce the loss of CTCF binding activity. Intriguingly, variants with the highest impact on binding activity had higher MAPS scores than missense variants. While the role of protein alterations in disease etiologies is well-established, we are only beginning to understand the functional consequences of CBS disruption. CBSs have a critical function in regulating gene expression by orchestrating the genome’s 3-dimensional (3D) organization^37–39,52^. Functional studies have linked the disruption of CBSs to both rare and complex disease; structural variants (SVs) that delete a CBS have been linked to pathogenic gene expression underlying rare limb disorders^7,53,54^, while various cancer types are enriched for SNVs in CTCF binding motifs^55,56^. These studies are in line with our findings here and together suggest an important role for CBSs in driving phenotypic variation in human populations. Disease studies primarily leverage a gene-based paradigm^57^. The high level of constraint we observed relative to missense variants further emphasizes the importance of regulatory annotations in prioritizing subsets of functional variants for disease studies.

We note several important limitations in our findings. Our annotation approach requires the uniform processing and integration of a TF’s binding patterns through ChIP-Seq across hundreds or even thousands of biosamples. Currently, this depth of data exists only for a few TFs, like CTCF. We anticipate the ability to annotate more TFs and therefore more SNVs as the cCRE framework expands. Furthermore, our summary of these binding patterns into a metric of activity is an effective means to distinguish likely functional binding sequences, which is critical for interpreting genetic variants. However, it does not incorporate tissue-specific activity and may down-weight CBSs that are highly active in poorly profiled tissues. Additionally, we note the bias inherent in thresholding for high-quality motifs during scanning. It remains unclear as to whether this bias contributed to the identification of substantially more loss of binding SNVs in gnomAD or if there are important differences in the functional properties of gain of binding SNVs.

Here, we integrated *in vitro* measurement of TF binding activity with information content encoded by PWMs for the annotation of noncoding SNVs. Our approach effectively captured regulatory variants with functional signatures in two computationally tractable steps: binding activity and statistical testing on ΔPWM scores. While conceptually similar frameworks exist, we found no viable alternative for the annotation of over 1 million SNVs in gnomAD. For example, the contextual analysis of transcription factor occupancy (CATO) method was previously introduced to capture allelic binding *in vitro* by integrating ΔPWM with DNase footprints across diverse celltypes^58^. However, in its current form, the application of CATO scores on data of this size is not feasible and there is no direct integration with external regulatory datasets, such as ENCODE’s cCREs. We expect that the relative simplicity and low computational cost of our approach will be crucial in facilitating future, large-scale annotation efforts on modern WGS.

Success with annotating genic variants (e.g. SpliceAI^59^) demonstrates the value of hypothesis driven frameworks^60^; interpretation of their outputs provides clear direction for subsequent studies. Until recently, the level of abstraction present in noncoding variant prioritization approaches has been a necessary and invaluable tool for overcoming limitations in data and a noncoding genetic code. However, their complexity and ambiguous outputs can severely limit their utility in the interpretation of noncoding genetic variants. For example, follow-up analyses on scores output from CADD may not always provide a clear biological interpretation and can risk becoming circular by applying further regulatory annotations to contextualize an elevated score. The barriers imposed by data availability and the lack of a regulatory code will continue to diminish as 1) large-scale consortia work to generate ever larger genetic and multi-omics datasets^61–63^ and 2) individual contributors advance conceptual frameworks to annotate noncoding DNA. We note that our effort to adapt a gene-like framework for noncoding variants is at the early stage and will require significant follow-up to assess the confidence and validity in our scores. Nevertheless, our approach has massively constrained the search space for novel, deleterious noncoding SNVs while providing a clear means to interpret their effects.

In summary, we introduce an approach to annotating noncoding variants that capitalizes on both the large population genome-sequencing datasets and extensive gene regulatory annotations that now exist. We demonstrate a conceptually and computationally tractable strategy to synthesizing these data and in doing so, find evidence for purifying selection acting against the loss of CTCF binding in human populations. We anticipate this work will aid in prioritizing functional regulatory variants with implications for understanding the biology of diseases and traits

## Methods

### 3.1. Annotating rDHS with CTCF binding activity

We downloaded the CTCF Z-score signal matrix^1^ for all representative DNase hypersensitivity sites (rDHSs) from the ENCODE-SCREEN web portal. We subset this matrix to rDHS with significant DNase signal in at least one biosample, which corresponds to the complete set of human candidate cis-regulatory elements (cCREs, n=1,063,879). We masked all rDHS-biosample combinations with a raw CTCF binding signal of zero (z-score equal to -10). To generate the binding activity annotation, we summarized each rDHS’s CTCF signal distribution using Stouffer’s meta-analysis of Z-scores^64^. To do this, we implemented the equation *zi/N*, where Zi represents a given biosample’s Z-score and N is the total number of non-masked biosamples for that rDHS.

### 3.2. Annotating CTCF motifs with binding activity and definition of activity quantiles

We downloaded all reference sequence matches to CTCF’s canonical motif (MA0139.2) in build hg38 from the JASPAR database. We then excluded sequence matches from non-autosomal and sex contigs, reported assembly gaps, ENCODE blacklisted regions^65^ and protein-coding exons (Gencode v44) using Bedtools *intersect*^66^. Our final track consisted of 1,764,648 sequence matches. To integrate this data with rDHS regions from ENCODE (above), we intersected the subset of rDHS that are cCREs with our final set of motifs using Bedtools *intersect*. We assigned each motif an activity score by reporting the activity of its intersecting cCRE. Motifs that did not intersect a cCRE were not assigned activity scores. In certain analyses, we refer to binding activity in terms of quantiles. To do this, we assigned the distribution of activity scores for all annotated rDHS into 100 equal sized bins.

### 3.3. Comparing predicted CTCF binding affinity with activity

We reported each CTCF motif’s predicted binding affinity as its precision-weight matrix (PWM) score, relative to the minimum and maximum score obtainable as in Gheorghe et al. 2019^67^. To characterize the relationship between the binding activity and these relative PWM scores, we binned each motif by their activity quantile (above) and assessed the mean relative PWM score in each bin. Their statistical evaluation was computed using a Pearson correlation.

### 3.4. Comparing evolutionary sequence conservation with CTCF binding activity

We assessed nucleotide conservation using four separate metrics. PhyloP100, PhastCons100 and GERP++ scores were downloaded in Bigwig format from the UCSC genome browser^68^. LINSIGHT scores were downloaded from Caleb et al. 2019^69^. We assigned all positions corresponding to motifs with rDHS support conservation scores using the package pyBigWig^70^. In the case of conservation scores generated in build hg19 (GERP ++ and LINSIGHT), we first lifted back each position’s coordinate using the UCSC LiftOver tool^71^. To assess the relative enrichment of conserved nucleotides against binding activity, we calculated the proportion of conserved nucleotides in each activity decile. We used thresholds of 2, 0.8, 0.8 and 1 to define conserved positions for GERP++, LINSIGHT, PhastCons100 and Phylop100, respectively. To assess confidence on each calculation, we randomly sampled with replacement nucleotides from each activity decile 10,000 times over 10 iterations. We plot the mean of these computations with their corresponding 95% confidence interval. We assessed the relationship and statistical significance using Spearman Correlation.

### 3.5. Processing genetic data from gnomAD

We downloaded the gnomAD v3.1.2 database and reduced it to SNVS, removing entries with allele count (AC) and frequency (AF) reported as zero. We further required all SNVs to be of high quality (QC=PASS) and be covered in at least half of the samples in gnomAD v3 (allele number, AN > 76,000). In total, we identified 569,860,911 SNVs after quality control. For each variant, we report its functional class as its most severe consequence term as assessed by the Variant Effect Predictor (VEP)^72^.

### 3.6. Annotating gain and loss of binding for all candidate SNVs in gnomAD

First, we identified all QC’d SNVs that overlap the subset of CTCF binding motifs with cCRE support using Bedtools *intersect*. To calculate ΔPWM scores and accompanying p-values, we used the R package atSNP^47^. To do this, we extracted 14bp of flanking reference sequence upstream and downstream of each variant’s genomic coordinate. We then report the highest log-odds score for the subsequence overlapping the variant’s position for both the REF and ALT alleles assessed by atSNP. Raw ΔPWM scores were quantified as the difference between the REF and ALT PWM score. We classify variants as putative gain of binding if their raw ΔPWM was <= 0 and loss of binding if their raw ΔPWM was > 0. To assess the significance of each observed ΔPWM score, we reported its –log10 p-value rank score, which is derived from atSNP’s importance sampling algorithm. To address truncation errors related to low p-values (p-value ∼ 0) we encoded dummy values of (10^-5).

### 3.7. Assessing constraint using scaled-CADD scores

We extracted PHRED-scaled CADD scores for each variant from the gnomAD v.3.1.2 hail table. To assess the relationship between these scores, binding activity and ΔPWM, we first stratified all variants by their class (gain of binding or loss of binding) and then by their significance (p-value < 0.05). For each binding activity decile, we randomly sampled variants with replacement 10000 times over 10 iterations. For each iteration, we calculated the proportion of variants that are putatively pathogenic using a scaled-CADD cutoff of 10. We plot the mean of these calculations and 95% confidence intervals.

### 3.8. Implementing MAPS scores

We implemented mutability adjusted proportion of singletons (MAPS) scores as described in Karzewski et al. 2020^11^ using custom scripts as follows. To annotate all SNVs in gnomAD with their trinucleotide mutation rate, we first identified their trinucleotide context by reporting each nucleotide upstream and downstream of the variant’s position using the package pySAM. Next, we downloaded trinucleotide mutation rate data from the gnomAD v2 flagship paper and merged it with each variant’s trinucleotide context. To calibrate the MAPS model, we used all QC’d SNVs with their most severe VEP annotation as “synonymous_variant” (n=2,161,831). We then referred to this trained model to predict MAPS for all other functional variant classes. For a given class, we calculated a raw singleton proportion as the number of singletons (AC=1) divided by the number of variants. We then applied the calibrated model to regress out the expected number of singletons within that bin given its average mutation rate. To assess confidence, we computed the standard error of the mean (SEM) for each bin.

