## Supplementary Material for "Identifying deleterious noncoding variation through gain and loss of CTCF binding activity"

Supplementary Table 1

| Annotation | N | Length (bp) | Footprint (Mb) | Coverage Conservation | Coverage CADD | Number SNVs |
|---|---|---|---|---|---|---|
| **rDHS** | 1063878 | 273 | 290 | -- | -- | -- |
| **rDHS + PWM** | 355418 | 15 | 5 | 100 | 0.24 | 1253330 |

**Supplementary Table 1. Summary statistics of the study dataset**. Statistics describing the noncoding sequence footprint of putative CBSs identified through our approach, stratified by whether the CBS has both rDHS and PWM support. The fields are as follows; N is the number of sequence elements in each group, Length is the mean length of all sequences for a given group in basepairs, footprint is the cumulative amount of genomic sequence covered through each annotation approach, coverage conservation describes the proportion of sequence annotated with evolutionary sequence conservation (GERP++, LINSIGHT, PhyloP100 and Phastcons100), Coverage CADD is the proportion of sequence annotated with CADD scores and Number of SNVs refers to the total number of overlapping SNVs in gnomAD. Dashed lines (--) signal data we did not collect as a part of this study.

| Supplementary Table 2 | | | |
|---|---|---|---|
| Variant Class | N | MAPS | SEM |
| Synonymous | 2926343 | 0.0000 | 0.0003 |
| 3'UTR | 10000000 | 0.0275 | 0.0002 |
| Intergenic | 19999998 | 0.0286 | 0.0001 |
| Missense | 6075537 | 0.0435 | 0.0002 |
| 5'UTR | 9442976 | 0.0558 | 0.0002 |
| Splice Donor | 112806 | 0.0965 | 0.0015 |
| Start Lost | 26198 | 0.0985 | 0.0030 |
| Stop Lost | 12784 | 0.1122 | 0.0043 |
| Stop Gained | 201570 | 0.1196 | 0.0011 |
| Splice Acceptor | 82838 | 0.1256 | 0.0017 |

**Supplementary Table 2.** MAPS scores for different functional classes of genic variation in gnomAD. Error was calculated as the standard error of the mean for each proportion. Variant class indicates the Variant Effect Predictor (VEP) worst consequence. The count of 3'UTR and Intergenic variants were limited to a random sample of the displayed size to facilitate their processing.

### Supplementary Figures

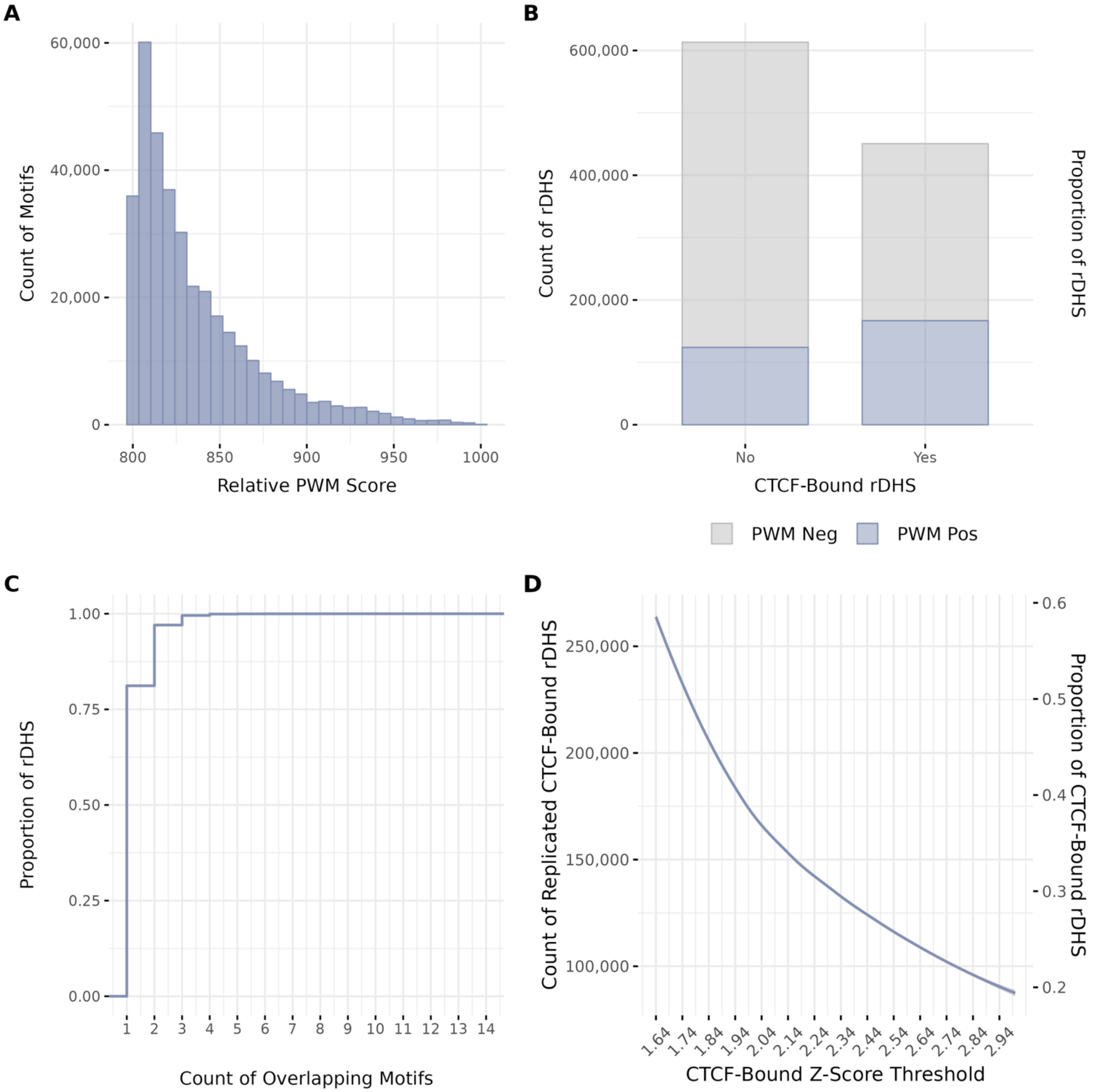


**Supplementary Figure 1. PWMs and ChIP-Seq alone are insufficient for the robust detection of active CTCF binding sequence. A)** Distribution of binding energies for all high-quality sequence matches (N = 1,764,648) to the canonical CTCF PWM (JASPAR MA0139.2) in the hg38 reference genome. The threshold for detection was a PWM score of 80% or higher relative to the max possible score for the CTCF PWM. **B)** Counts of rDHS stratified by whether the rDHS is classified as “CTCF Bound“ using criteria from ENOCODE (Z-score > 1.64 in at least biosample). Each group is colored to reflect the proportion of rDHS that have >=1 overlapping CTCF binding motif. **C)** Cumulative proportion of rDHS by the number of overlapping CTCF motifs after reducing to rDHS with >=1 overlapping motif. **D)** Count of rDHS called as CTCF bound cCREs by ENCODE that are replicated in >= 1 biosample at varying detection thresholds. The right Y-axis measures the ratio of replicated CTCF bound rDHS at a given threshold to the total number of cCREs called as CTCF bound (n=450,641).

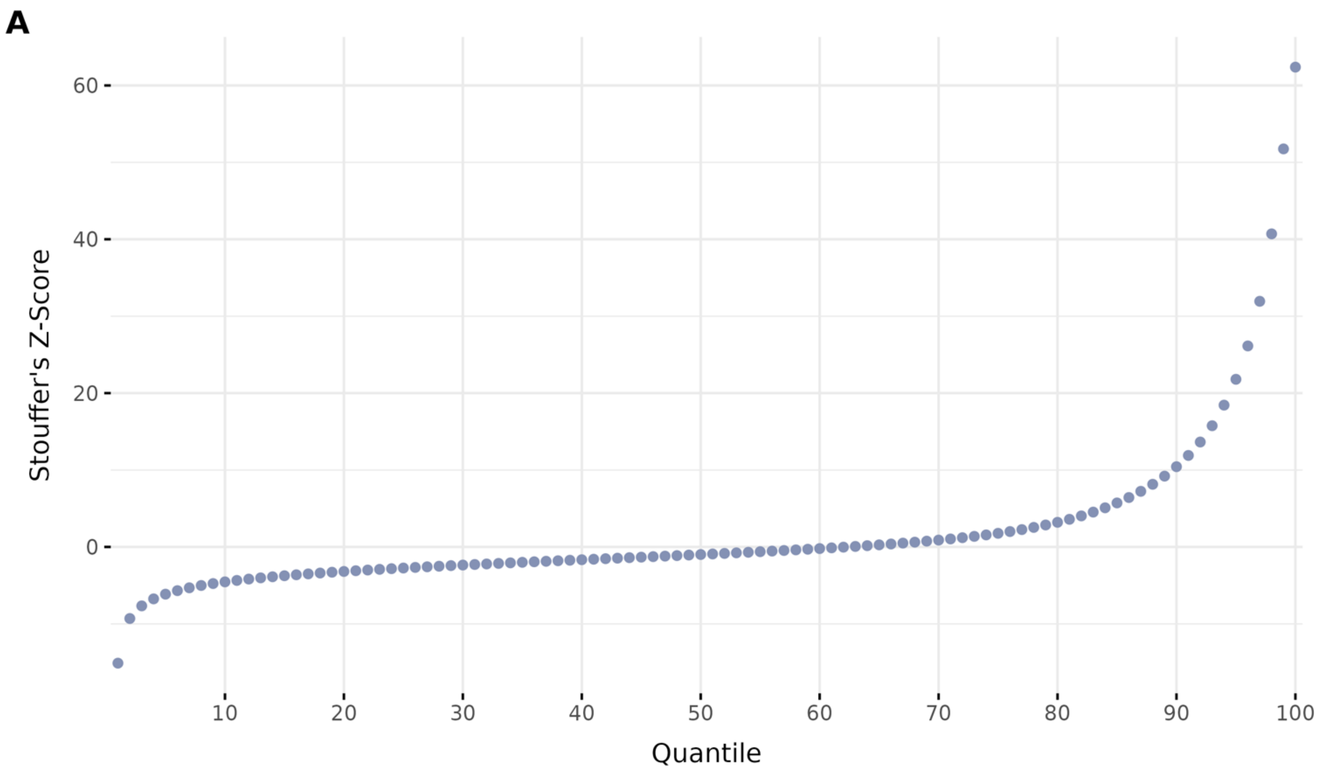


**Supplementary Figure 2. Overview of meta-analyzed Z-scores for each quantile of binding activity.** Distributions of meta-analyzed binding activity scores after binning all rDHS into 100 equal sized bins. Each quantile contains approximately 10,638 unique rDHS.

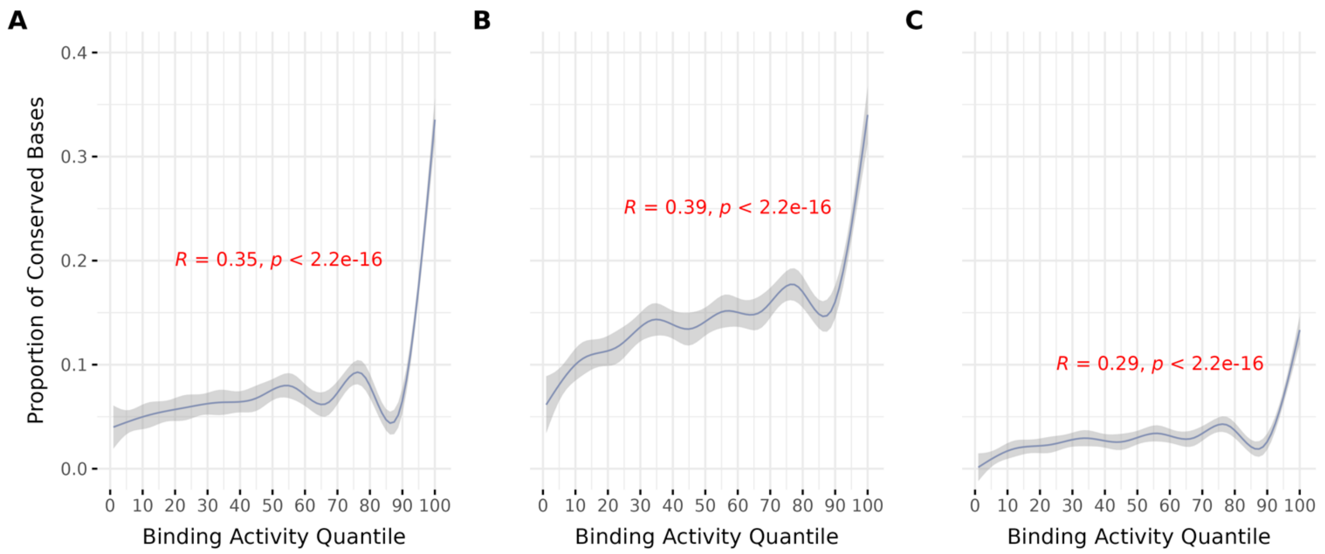


**Supplementary Figure 3. The enrichment of evolutionary sequence conservation at high binding activity quantiles is robust across different conservation metrics.** Relationship between evolutionary sequence conservation of CTCF motifs and binding activity quantiles. Conservation was measured as a proportion of conserved CTCF motif positions within each quantile using GERP++, LINSIGHT and Phastcons100 scores. Conservation was measured as a proportion of conserved bases within each bin using a threshold of 2, 0.8 and 0.8 for GERP++, LINSIGHT and Phastcons100 respectively.

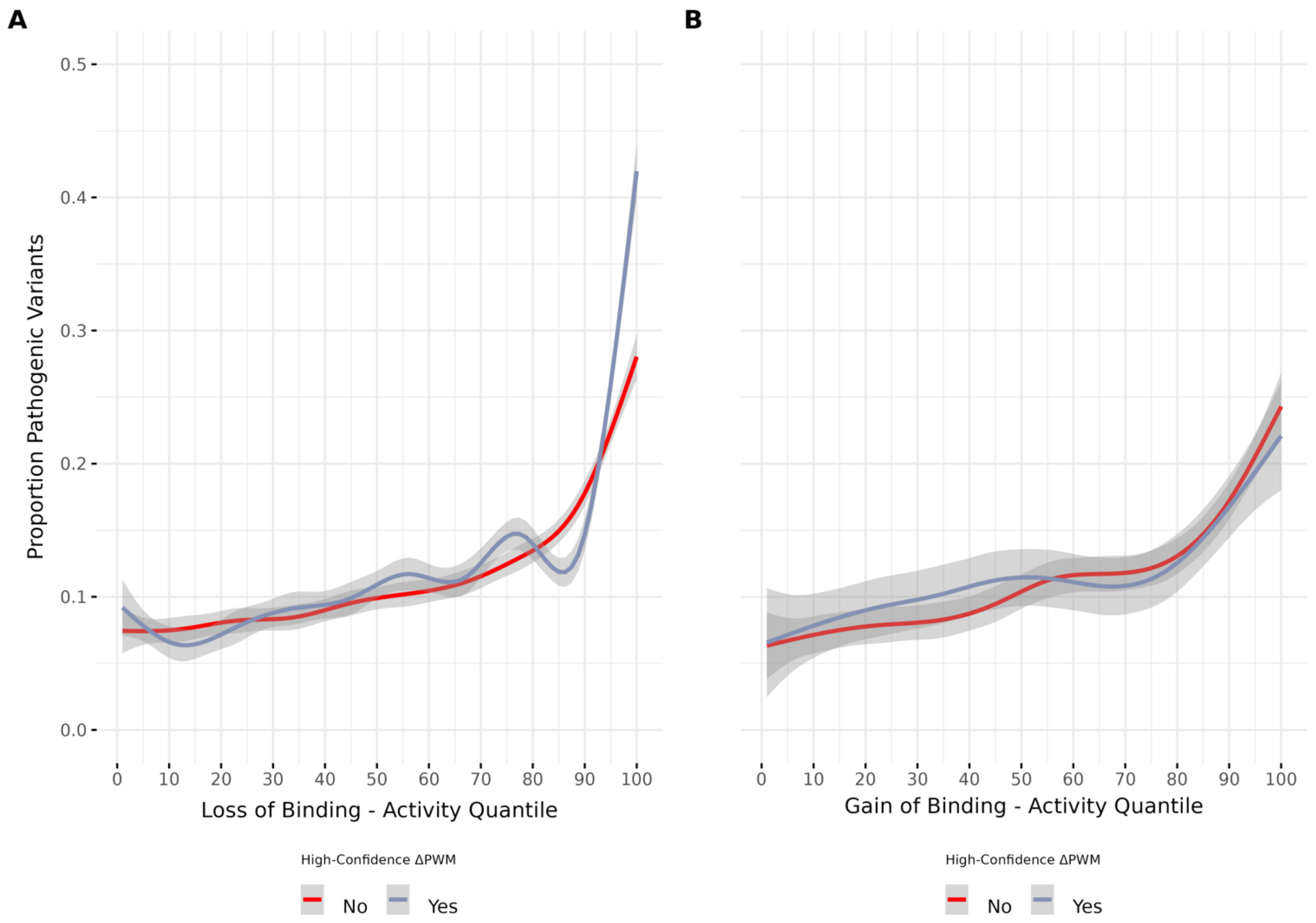


**Supplementary Figure 4. A)** Relationship between proportion of putative pathogenic SNVs based on scaled CADD scores and the loss of CTCF binding activity. A scaled CADD score of >=10 was used as a threshold of pathogenic or not. Y-axis displays the proportion of pathogenic SNVs within each activity quantile. Error as a 95% bootstrapped confidence interval. Colors indicate stratification on the confidence of the ΔPWM call using a threshold of 0.05. **B)** Relationship between proportion of putative pathogenic SNVs based on scaled CADD scores and the gain of CTCF binding activity assessed using the same approach as in A.

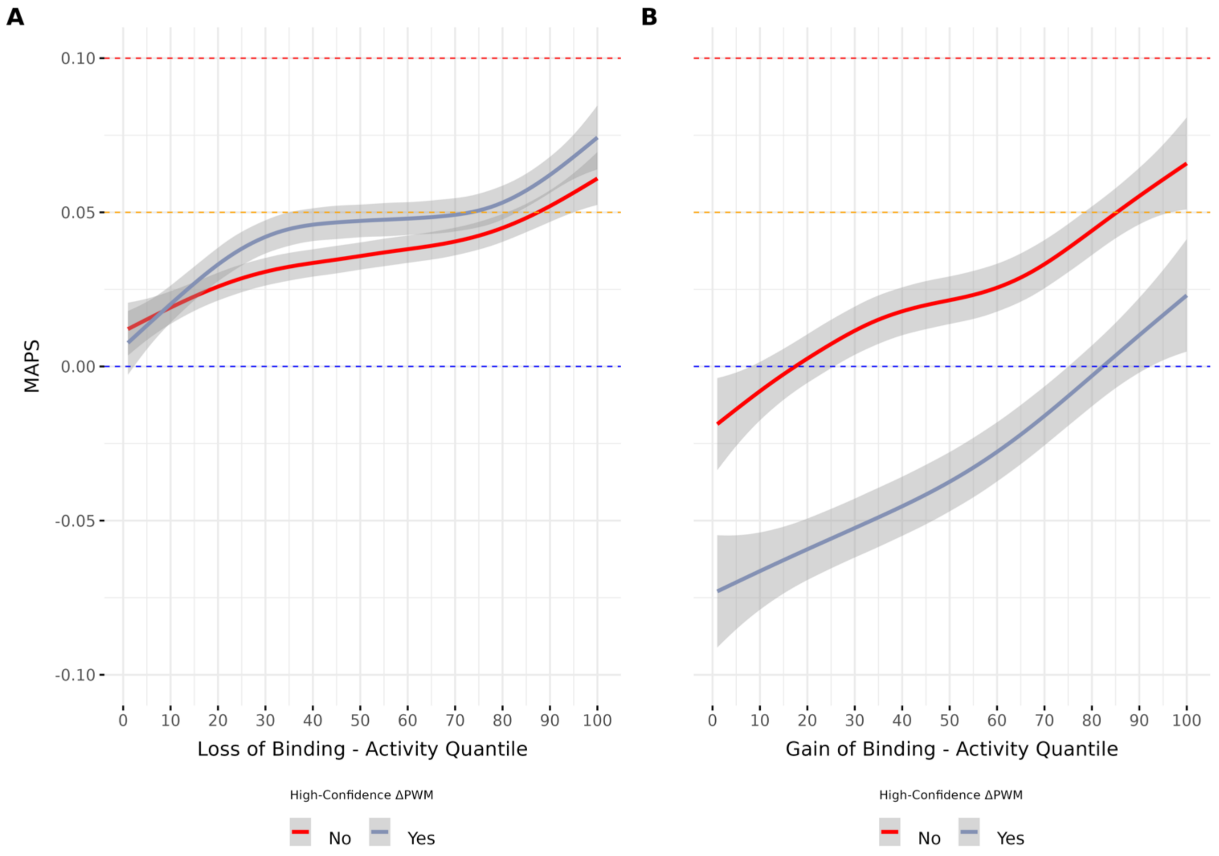


**Supplementary Figure 5. A)** Relationship between allele frequence as measured by MAPS and the loss of CTCF binding activity stratified by confidence in the ΔPWM calls. For context, we display MAPS scores for synonymous (blue line), missense (orange) and splicing (red) variants. **B)** Relationship between allele frequence as measured by MAPS and the gain of CTCF binding activity stratified by confidence in the ΔPWM calls.

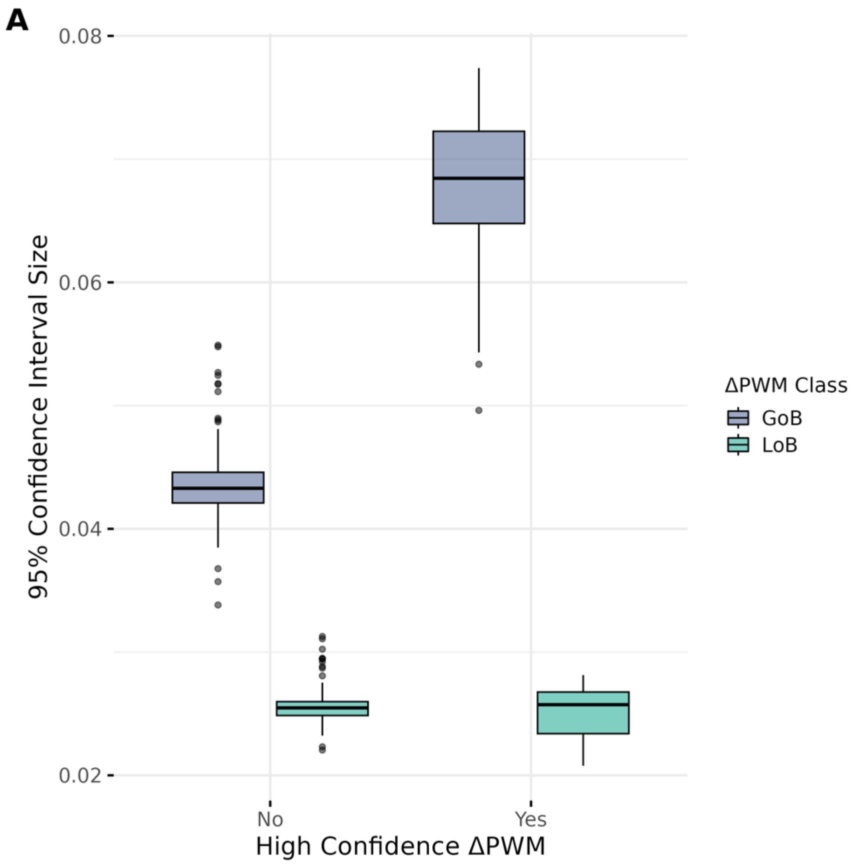


**Supplementary Figure 6.** Size of 95% confidence intervals on all singleton proportions used to calculate MAPS scores. Confidence was measured by applying a binomial test on the proportion of singletons variants in each binding activity quantile, stratified by the confidence in each variant's ΔPWM score. Gain of binding and loss of binding are abbreviated as GoB and LoB, respectively.
